# Worldwide population structure, long term demography, and local adaptation of *Helicobacter pylori*

**DOI:** 10.1101/019430

**Authors:** Valeria Montano, Xavier Didelot, Matthieu Foll, Bodo Linz, Richard Reinhardt, Sebastian Suerbaum, Yoshan Moodley, Jeffrey D. Jensen

## Abstract

*Helicobacter pylori* is an important human pathogen associated with serious gastric diseases. Owing to its medical importance and close relationship with its human host, understanding genomic patterns of global and local adaptation in *H. pylori* may be of particular significance for both clinical and evolutionary studies. Here we present the first such whole-genome analysis of 60 globally distributed strains, from which we inferred worldwide population structure and demographic history and shed light on interesting global and local events of positive selection, with particular emphasis on the evolution of San-associated lineages. Our results indicate a more ancient origin for the association of humans and *H. pylori* than previously thought. We identify several important perspectives for future clinical research on candidate selected regions that include both previously characterized genes (*e.g.* transcription elongation factor *NusA* and tumor Necrosis Factor Alpha-Inducing Protein *Tipα*) and hitherto unknown functional genes.

## Introduction

*Helicobacter pylori* is a Gram-negative bacterium that infects the mucosa of the human stomach. It was first described in the 1980s, when it was initially identified in association with chronic gastritis and later causally linked to serious gastric pathologies such as gastric cancer and ulcers (Marshall and Warren 1984; Suerbaum and Michetti 2002). It infects more than 80% of humans in developing countries and, although its prevalence is lower in developed countries, nearly 50% of the worldwide human population is infected (Ghose et al. 2005; Salih 2009; Salama et al. 2013).

Due to its clinical and evolutionary importance, there has been considerable research on mechanisms of *H. pylori* transmission, as well as on the population genetics and phylogenetic relationships among global isolates. Thus far, population genetic analyses have mainly focused on seven housekeeping genes (usually referred to as MLST), with the primary conclusions being that *H. pylori* strains appear highly structured, and their phylogeographic patterns correlate consistently with that of their human hosts. Given that the *H. pylori* -humans association is at least 100 kya old (Moodley et al. 2012), the current population structure of *H. pylori* may be regarded as mirroring past human expansions and migrations (Falush et al. 2003; Linz et al. 2007; Moodley and Linz 2009; Breurec et al. 2011) and thus help us shed light on yet unknown dynamics of local demographic processes in human evolution. However, despite the knowledge gained thus far, the long-term global demographic history of *H. pylori* has never been directly inferred.

The long, intimate association of *H. pylori* with humans suggests a history of bacterial adaptation. Considerable attention has focused on specific genes involved in modulating adaptive immunity of the host (for a review see Yamaoka 2010 and Salama et al. 2013) and on genomic changes occurring during acute and chronic *H. pylori* infection (Kennemann et al. 2011; Linz et al. 2014) as well as during *H. pylori* transmission between human hosts (Linz et al. 2013). However, bacterial genome adaptation has not been investigated at the global level. Owing to the recent introduction of next generation sequencing approaches, several complete *H. pylori* genomes have been characterized and are now available to further explore the selective history that might have contributed to shaping the bacterial genome.

Here, we study a combined sample of 60 complete *H. pylori* genome sequences (53 previously published, 7 newly sequenced) with origins spanning all five continents. Our aims were to detect adaptive traits that are commonly shared among the worldwide *H. pylori* population as well as to uncover patterns of local adaptation. We expect that, apart from a generally important role of adaptation to the human gastrointestinal environment, the differing eco-physiological conditions found in the gastric niche of worldwide human hosts, based on diverse diets and different bacterial compositions, could likely generate differential selective pressure on specific bacterial traits leading to locally adaptive events. For instance, an increase in pathogenicity seems to have occurred in *H. pylori* during the colonization of East Asia and could be partially explained by the presence of different alleles of virulence factors (*e.g*. CagA, VacA and OipA; Yamaoka 2010); also, colonization of the stomach niche has been optimized by regulation of motility and by bacterial cell shape (Sycuro et al. 2012).

To disentangle the signatures of demographic processes from the effects of natural selection on the distribution of allele frequencies, we first investigated the demographic history of our worldwide genome sample. Given that the genetic structure retrieved among the bacterial genomes mirrors the geographic distribution of human populations (Moodley and Linz 2009; Breurec et al. 2011; Moodley et al. 2012), the vast literature on human demographic history provides a solid basis for the study (*e.g*., Cavalli-Sforza et al. 1994), but modelling human-*H. pylori* co-evolution would also require knowledge of transmission dynamics and within-host variation. Despite the large number of surveys carried out, *H. pylori* transmission via an external source has never been demonstrated and direct contact among individuals is still considered the predominant mechanism (Brown 2000; Van Duynhoven and De Jonge 2001; Allaker et al. 2002; Perry et al. 2006). Transmission also depends on the hosts’ access to health care and socio-economic conditions. In developing countries, *H. pylori* transmission seems to happen preferentially but not exclusively among individuals who are closely related or living together (Schwarz et al. 2008; Didelot et al. 2013). However, in developed countries, improved hygienic conditions have decreased *H. pylori* prevalence, and transmission occurs primarily between family members, especially from mothers to children (Bureš et al. 2006; Chen et al. 2007; Khalifa et al. 2010; Krebes et al. 2014). Further, an important epidemiological factor is that a human host is normally infected with *H. pylori* within the first five years of life and, unless treated, infection persists the entire host lifespan. The host individual is therefore always potentially infective.

The human stomach is typically infected with a single dominant strain, with multiple infections occurring less frequently (*e.g*. Schwarz et al. 2008; Morelli et al. 2010; Nell et al. 2013). However, this empirical observation may be due to an experimental approach that intrinsically limits the detection of multiple infections (Didelot et al. 2013) since only a single isolate per patient is generally studied, and more focused approaches have highlighted higher within host variation (Ghose et al. 2005; Patra et al. 2012). In addition, MLST studies have detected a small fraction of human hosts from the same population sharing the same bacterial strain (or at least highly related strains with identical sequence type) (Patra et al. 2012; Nell et al. 2013). At the molecular level, mutation and recombination have been identified as the key forces responsible for population genetic variability (Suerbaum and Josenhans 2007). A recent whole genome study on 45 infected South Africans demonstrated that recombination is the major driver of diversification in most (but not all) hosts (Didelot et al. 2013), confirming previous observations (Falush et al. 2001; Kennemann et al. 2011). At the population level, recombination is very frequent throughout the genome along with other events such as rearrangements, transpositions, insertions and gene gain or loss (Gressmann et al. 2005; Kawai et al. 2011). The relative roles of demographic and selective processes in shaping the bacterial genetic variation during the lifespan of a single host have yet to be explored.

Given our limited knowledge of *H. pylori* epidemiology and thus its consequences on long-term evolution, we here explore the species’ genetic structure using newly available worldwide genomic data to infer the demographic history of the sampled populations, directly addressing the extent to which the population history of *H. pylori* mirrors that of its human host. Using this estimated demographic model as a null, we explore two different approaches in order to characterize both local and global events of positive selection. Our results indicate global signatures of selection in functionally and medically relevant genes and highlight strong selective pressures differentiating African and non-African populations, with over one hundred putatively positively selected genes identified.

## Materials and Methods

### Samples and whole genome sequencing

Seven complete *H. pylori* genomes were newly typed for the present study to increase the currently available set of 53 genomes, in order to represent all five continents (Table S1). The most valuable contributions among our sequences were the Australian aboriginal, Papua New Guinean Highlander, Sudanese Nilo-Saharan and South African San genomes, which have never been previously characterized.

Data production was performed on a ROCHE 454 FLX Titanium sequencer. WGS sequencing libraries for pyrosequencing were constructed according to the manufacturers’ protocols (Roche 2009 version). Single-end reads from 454 libraries were filtered for duplicates (gsMapper v2.3, Roche) and could be directly converted to frg format that was used in the genome assembler by Celera Assembler v6.1 (CA6.1; Miller et al. 2008). Several software solutions for WGS assembly were tested during the project among them Roche’s Newbler and CeleraAssembler (both can assemble all read types). Genome assembly was performed on a Linux server with several TB disk space, 48 CPU cores and 512 GB RAM.

### Bioinformatics

After long read assembly, the seven new genomes were further re-ordered using the algorithm for moving contigs implemented in Mauve 2.3.1 software for bacterial genome alignment (Darling et al. 2004; 2010). In this analysis, the scaffold sequence for each genome to be reconstructed was assigned on the basis of geographical proximity. In particular, the sequences from Papua New Guinea and Australia were re-ordered against an Indian reference (*Helicobacter pylori* India7, GenBank reference: CP002331.1; see Table S1), given the absence of closer individuals. The global alignment of the genomes was carried out using mauveAligner in Mauve 2.3.1 with seed size calibrated to ∼12 for our data set (average size ∼1.62 megabases). The minimum weight for local collinear blocks, deduced after trial runs performed using default parameter settings, was set to 100. The original Mauve alignment algorithm was preferred to the alternative progressive approach (progressiveMauve; Darling et al. 2010) because of its higher performance among closely related bacterial genomes (appropriate in the present case of intraspecific analysis), its higher computational speed, and to avoid the circularity of estimating a guide phylogenetic tree to infer the alignment. The aligned sequences shared by all genomes were uploaded into R using the package *ape* (Paradis et al. 2004) and processed for post alignment refinement. The length of the genomes prior to alignment ranged from 1,510,564 bp to 1,709,911 bp with an average of 1,623,888 bp. The Mauve alignment consisted of 71 blocks commonly shared by all the individuals for a total of 2,586,916 sites. Loci with more than 5% missing data were removed, giving a final alignment of length to 1,271,723 sites. The final number of segregating sites in the global sample was 342,574 (26.9%). Among these, we found 302,278 biallelic sites and 35,003 and 5,293 tri- and tetra-allelic sites respectively. The distribution of segregating sites along the aligned sequences is shown in Figure S1.

### Structure analysis

In order to first define the populations to be used in subsequent analyses, we compared a multivariate approach, discriminant analysis of principal components (DAPC; Jombart et al. 2010) with two Bayesian analyses of population structure BAPSv5.4 (Corander et al. 2006, 2008) and STRCTURE (Pritchard et al. 2000). The first method assesses the best number of clusters optimizing the between-and within-group variance of allele frequencies and does not assume an explicit biological model, while the second is based on a biological model that can also detect admixture among individuals. The optimal number of population clusters was established by both methods. In DAPC this is done through the Bayesian information criterion (BIC) using the *find.clusters* function in *adegenet* 3.1.9 (Jombart 2008; Jombart and Ahmed 2011) while BAPS estimates the best *K* comparing the likelihood of each given structure. We ran the DAPC analysis with 1,000 starting points and 1,000,000 iterations and found that results were consistently convergent over 10 independent trials. BAPS was run with a subset of 100,000 SNPs using the admixture model for haploid individuals and was shown to be effective to detect bacterial populations and gene flow in large-scale datasets (Tang et al. 2009; Willems et al. 2012). STRUCTURE was run on a subset of 100 kb, for a total of 29,242 SNPs, using 10,000 burn-in and 50,000 iterations, and we replicated 5 runs for each tested number of partitions (from 2 to 10) with the admixture model. Finally, the seven housekeeping genes historically used in *H. pylori* population genetics (MLST) were extracted from the alignment and used to assign populations to strains with STRUCTUREv2.3.4 (Falush et al. 2007) as a comparison with previous work.

For further insight into population structure we reconstructed the clonal genealogy of bacterial genomes using ClonalFrame v1.2 (Didelot and Falush 2007). This method reconstructs the most likely clonal genealogy among the sequences under a coalescent model with mutation and recombination, so that the model of molecular evolution takes into account both the effect of mutated sites and imported (recombining) sites. We also evaluated fine-scale population structure from sequence co-ancestry using fineSTRUCTURE (Lawson et al. 2012). This method performs Bayesian clustering on dense sequencing data and produces a matrix of the individual co-ancestry. Each individual is assumed to “copy” its genetic material from all other individuals in the sample, and the matrix of co-ancestry represents how much each individual copied from all others.

Population summary statistics (the number of segregating sites, genetic diversity, mean number of pairwise differences, Tajima's *D*, and pairwise *F*_*ST*_) were estimated with R packages adegenet and pegas (Paradis 2010).

### Inferring demographic history

The genomic landscape is shaped by the combined evolutionary signature of population demography and selection. Not accounting for population demography, therefore, could lead to biased estimates of both the frequency and strength of genomic selection (*e.g.*, Thornton and Jensen 2007).

While many of the available statistical methods for detecting patterns of genome-wide selection have been argued to be robust to demographic models of population divergence and expansion (Nielsen et al. 2005; Jensen et al. 2007b; Foll and Gaggiotti 2008; Narum and Hess 2011), they also have limitations (Narum and Hess 2011; Crisci et al. 2013). In highly recombining species such as *H. pylori* (Morelli et al. 2010; Didelot et al. 2013), evidence of recent positive selection events across the global population may have become obscured, owing to the reduced footprint of selection.

It was therefore necessary to first explicitly infer the demographic history, in order to disentangle the effects of population demography on the allele frequency distribution from the possible effects of selective processes. Here, we tested different neutral demographic scenarios, making assumptions based on the observed genetic structure and previous knowledge of human evolutionary history.

Demographic scenarios were modelled and implemented in the software *fastsimcoal2.1* (Excoffier et al. 2013), allowing for the estimation of demographic parameters based on the joint site frequency spectrum of multiple populations. The software calculates the maximum likelihood of a set of demographic parameters given the probability of observing a certain site frequency spectrum derived under a specified demographic model. This program uses non-binding initial search ranges that allow the most likely parameter estimates to grow up to 30%, even outside the given initial search range, after each cycle. This feature reduces the dependence of the best parameter estimates on the assumed initial parameter ranges. Model details and initial parameter range distributions are given in Supplementary Materials (see Files S1 and S2). We assumed a finite site mutation model, meaning that the observed and simulated joint site frequency spectra were calculated to include all derived alleles in multiple hit loci (Figure S6).

### Model choice and demographic estimates

Firstly, different tree topologies based on hierarchical structures, as obtained with the approaches described above, were compared to infer the best population tree, assuming divergence without migration. Once the tree topology with the strongest statistical support was established, we evaluated and compared the likelihood of models including asymmetric migration among populations. Migration models were tested starting with interchanging individuals only among single pairs of closely related populations. We could therefore assess whether adding migration would improve the likelihood compared to a divergence-without-migration model, and which pairs of populations are most likely to exchange migrants. We also allowed migration among more distantly related populations in addition to a simple pairwise stepping stone model.

The best model among those tested was selected through the corrected Akaike Information criterion (AICc) based on the maximum likelihoods calculated for independent runs.

### Testing models of positive selection

Two different statistical tests were used to detect global and local candidate loci for selection. First we used the SweeD algorithm (Pavlidis et al. 2013), derived from SweepFinder (Nielsen et al. 2005) to localize recent events of positive selection, an approach based upon comparison with the ‘background’ site frequency spectrum (Figure S7). The scan for positive selection is carried out by centering the maximized probability of a selective sweep on a sliding-window locus along the chromosome, and calculating the composite likelihood for each centered locus to fall within a region where the distribution of SNPs deviates from the neutral expectation. When an outgroup sequence is available to establish derived mutations, the empirical site frequency spectrum estimated from the observed dataset is unfolded, otherwise only minor alleles are used for the calculation (*i.e*., a folded SFS). Given the difficulties associated with bacterial genome alignment of suitably close outgroup species, we ran our estimates on a folded SFS. All tri- and tetra-allelic SNPs were removed, and monomorphic loci were not considered in the calculation and the grid was set to 500,000 bp. We analyzed the entire dataset (60 genomes) as well as each of the five populations separately.

Second, we applied a method based on the detection of patterns of linkage disequilibrium (LD) around a SNP (OmegaPlus; Kim and Nielsen 2004; Jensen et al. 2007a; Alachiotis et al. 2012), since LD is expected to result from a selective sweep owing to the hitchhiking of linked neutral mutations (Maynard Smith and Haigh 1974). This complements the SFS approach as it is applicable to sub-genomic regions, contrary to SweeD, and it has proven effective under specific demographic models for which SFS-based approaches are less powerful (Jensen et al. 2007a; Crisci et al. 2013). We used windows of size between 1,000 and 100,000 base pairs.

Finally, a total of 1000 simulated data sets, generated using most likely demographic parameter estimates, were analyzed with SweeD and OmegaPlus in order to gain an empirical distribution of likelihoods (SweeD) and omega values (OmegaPlus) in a neutrally evolving population. The only parameter drawn from a range was the recombination rate, calibrated around the most likely estimate obtained with ClonalFrame, with the aim of providing an empirical evaluation of its impact on the methods we used to infer selection. The simulated distribution of these selection statistics, based upon the previously inferred demographic history, allows for statistical statements to be made regarding the likelihood that observed outliers are consistent with neutrality alone. A *p*-value for each observed omega and likelihood was obtained using the function *as.randtest* of *ade4* R package, calculated as (number of simulated values equal to or greater than the observed one + 1)/(number of simulated values + 1).

### Gene annotation and biological interpretation of the results

Annotation of the bacterial genes was performed using the free automated web server *BASys* (Bacterial Annotation System, www.basys.ca; Van Domselaar et al. 2005). The annotation was run on aligned sequences, removing multiply hit loci. The annotated genome of Africa1 is provided as an example in Supplementary File 3, and all annotation files are available upon request. The regions identified as being under selection were then compared with the gene annotation.

## Results

### Population structure and genetic diversity

Given the difficulties of defining a population among a bacterial sample, we decided to perform our cluster analysis using three approaches (DAPC, BAPS and STRUCTURE) that rely on very different assumptions, keeping in mind that using semi or fully parametric methods (such as STRUCTURE-like approaches) is more likely to lead to violation of the methodological assumptions and therefore to biased results (Lawson 2013). DAPC may out-perform STRUCTURE when dealing with data sets with a high degree of isolation by distance (e.g. Kalinowsky 2011), as it is likely the case for *H. pylori* populations (Linz et al. 2007; Moodley and Linz 2009), and it also provides the possibility of visualizing clusters' reciprocal distances in the multivariate discriminant space. BAPS and STRUCTURE, on the other hand, offer a biological model to test individual admixture, which is particularly useful to gain an understanding of the degree of differentiation, such that these methodologies may be considered complementary_._ Population structure analyses were consistent between the model-free DAPC and model-based BAPS and STRUCTURE approaches. All structure approaches were in agreement on a worldwide number of populations that does not exceed *K* = 4.

DAPC indicated *K* = 4 as the best clustering (Figure S2A) while BAPS estimates *K* = 3 and STRUCTURE analysis offers a best *K* in between 2 and 4, with most support for *K* = 3 and partitions above 5 dramatically decreasing the likelihood (Figure S2B). Most importantly, the three methods are in consistent agreement on the assignment of single individuals to clusters (Table S1). With the least hierarchical division (*K* = 3), one population comprised African genomes containing all strains from Khoisan-speaking human hosts (referred to as Africa2; Figure 1A and Figure S3). Other African and European strains fell into a population cluster, called here AfricaEu (Figure 1A and Figure S3). A final population is composed of Central Asian, Sahul, East Asian and Amerind strains (AsiaAmerica; Figure 1A and Figure S3). Finer structuring (*K* = 4) separates the non-Khoisan African sequences (Africa2 and Africa 1), but merged European with Central Asian sequences into a new population (referred to as EuroAsia), with Asian and American strains making up the fourth cluster (AsiaAmeria). The only difference between DAPC, BAPS and STRUCTURE analyses at *K* = 4 is given by individual 7, which is clustered in the AsiaAmerican or EuroAsian populations, respectively. At *K* > 4, American strains were separated into a fifth independent cluster by DAPC, but not by BAPS or STRUCTURE. Plotting the first two discriminant components (DCs) for *K* = 4 (Figures 1B) most strikingly depicted the second African cluster as highly divergent along DC1, whereas divergence among the other clusters was mainly along DC2.

**Figure 1.**
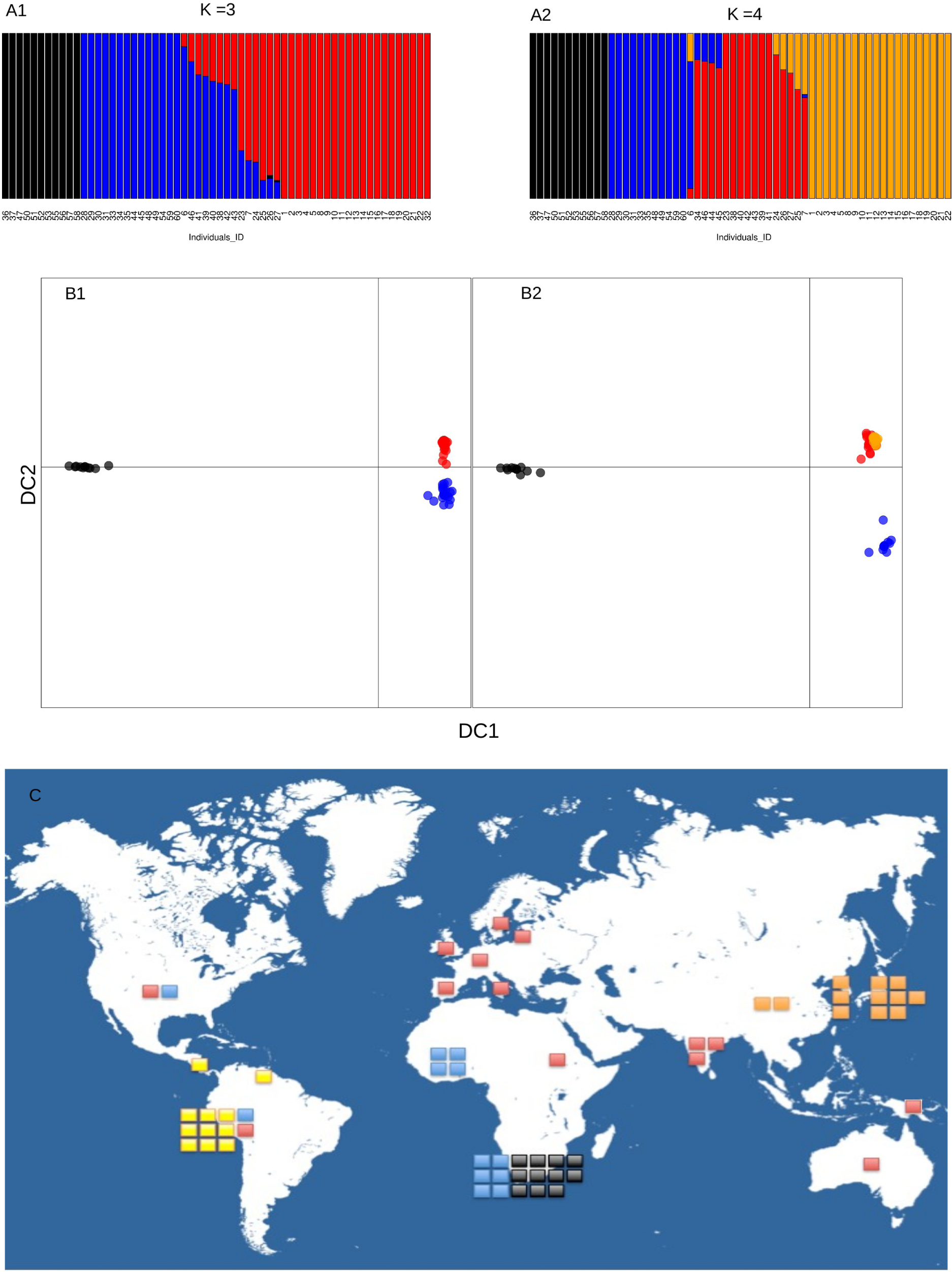
A) Plots of individual assignments to clusters according to BAPS and DAPC analysis, using *K* = 3 (A1) clusters with black = Africa2, blue = AfricaEu, red = EuroAsia; and *K* = 4 (A2) clusters with black = Africa2, blue = Africa1, red = EuroAsia and orange = AsiaAmerica; B). Scatterplots of the discriminant space (components 1 and 2), using *K* = 4 (B1), *K* = 5 (B2); C). World map with squares representing individuals colored according to cluster assignments with yellow squares indicating American sub-cluster (as for *K* = 5 in DAPC analysis; see STable 1).

The clonal genealogy (Figure S4) and analysis of fine structure (Figure S5) were in strong agreement with the geographical structuring elucidated by previous approaches. The Africa2 population was well differentiated in the genealogical tree (Figure S4) and in the co-ancestry matrix (Figure S5), while the remaining populations appear more closely related, and all non-African strains formed a clearly monophyletic clonal group. Asian and American populations were well differentiated in the co-ancestry analysis and were divided into distinct sub-clades in the clonal genealogy. The two Sahul genomes shared a higher degree of relatedness with three Indian genomes and these did not cluster monophyletically with the other Eurasian genomes in the clonal genealogy, instead clustering geographically between Eurasian and East Asian groups (see both Figure S4 and S5). Individual 7 appeared intermediately related to both Indian-Sahul and the more divergent Amerind strains. In the following analyses, this strain was left within the European population as indicated by BAPS and also by STRUCTURE analyses of the MLST data (hpEurope).

The population genomic structure elucidated here is in agreement with previous analyses of global structuring of MLST genes, where the highest diversity was found among African strains, the most divergent being the population hpAfrica2 (Falush et al. 2003). They also agree that Central Asian (hpAsia2), North-East African (hpNEAfrica) and European (hpEurope) strains are closely related (Linz et al. 2007) and sister to hpSahul (Australians and New Guineans, Moodley et al. 2009), and that East Asian and Amerind strains (hpEastAsia) share a relatively recent common ancestor (Moodley and Linz 2009). The divergent hpAfrica2 was shown to have originated in the San, a group of click-speaking hunter-gatherers whose extant distribution is restricted to southern Africa (Moodley et al. 2012). A complete list of individuals, geographic origin and cluster assignment based on DAPC, BAPS and STRUCTURE (100kb and MLST extracted from our alignment) is given Table S1. Predictably, genetic diversity indices were highest for the Eurasian population containing the geographically diverse strains from North East Africa-Europe-Central Asia and Sahul, especially evident from the number of tri-allelic and tetra-allelic loci and the mean number of pair-wise differences, while the Amerind population was most homogeneous (Table 1). It is worth noting that within the EuroAsian population there is the highest nucleotide diversity, as European sequences show a value of 0.042 (s.d. 0.0005), the three Indian strains 0.038 (± 0.0008) and the only two Sahul sequences 0.036 (±. 0.0013). Only the Africa1 population reaches such value of internal diversity (0.038 ± 0.0003), while all the others fell below 0.03.

**Table 1.** Population summary statistics based on a globally representative data set of 60 *Helicobacter pylori* genomes. *N* is the number of strains per population; *n* is the mean number of pairwise differences. P-values refer to Tajima's *D*.

| Pop | N | Number of segregating Loci |  |  | n | Tajima's D | p-value |
| --- | --- | --- | --- | --- | --- | --- | --- |
|  |  | 2 alleles | 3 alleles | 4 alleles |  |  |  |
| Africa2 | 11 | 117125 | 4276 | 171 | 40472.9 | -0.40437 | 0.747 |
| Africa1 | 12 | 139958 | 7034 | 345 | 50885.2 | -0.03660 | 0.995 |
| EurAsia | 16 | 197093 | 16713 | 1325 | 63187.7 | -0.52261 | 0.649 |
| EastAsia | 12 | 127160 | 5508 | 275 | 40246.74 | -0.79713 | 0.476 |
| America | 9 | 101895 | 3777 | 183 | 30781.8 | -0.47308 | 0.706 |

### Demographic inference

Overall, the different clustering methods and genealogical approaches implemented here were largely consistent in their population assignment. Although the American cluster appears to be more likely sub-structure, we included it into the further analyses as a separated population. This is owing to the fact that the demographic and selective history associated with the peopling of the Americas would suggest that this group of strains have likely undergone a very different fate than the East Asian strains with which they are closely related. This notion seems indeed to be confirmed by the population-specific tests of positive selection presented below. Furthermore, treating American strains separately offers the possibility of testing the hypothesis of a concerted bacterial-human expansion, as the timing of human colonization of the Americas is a well-characterized event, allowing for comparison with our inference. We proceeded hierarchically to test different genealogical topologies building on the population structure outlined above. First we tested the hypothesis of three main worldwide populations (*K* = 3, panel A, Fig. S5), with Africa2 strains forming the most ancestral population, in agreement with our and previous findings (Moodley et al. 2012). Alternative origins of the two other clusters – AfricaEu and AsiaAmerica – were therefore tested in three possible topologies (1-3, panel A, Fig. S5), with these two populations derived after an ancient split with the Africa2 ancestral population (Figure S6). A comparison of likelihoods suggests the first genealogical setting (see Figure S6) as the most supported, that is, AfricaEu strains are more ancestral than Eastern Asian and American strains, following a pattern close to that of human expansion (Table S2A).

Introducing a further population subdivision (i.e., *K* = 4), we tested different hypotheses for the origin and timing of the out-of-Africa sub-populations, that is EuroAsia and AsiaAmerica (Panel B, Figure S6). Lastly, we considered an additional sub-population formed by American strains, in agreement with DAPC subdivision at *K* > 4 (panel C, Figure S6). Clearly, the addition of multiple populations decreases the degrees of freedom and likelihood value of demographic models, and the hierarchical levels A, B and C are thus not directly comparable. However, in all tests, a model of population split resembling human expansion out-of-Africa was always preferred (Table S2A). The results of demographic inference for models without migration were highly compatible across different population sub-structures (Table S2A).

Finally, hierarchical models based on five populations, and using the most likely genealogical topology obtained with a purely divergent model, were also tested under the assumption of asymmetrical between lineage gene flow. Each time a pairwise asymmetric migration rate improved the likelihood of the model, the same scenario was re-analysed adding a further pairwise migration rate, for a total of 20 demographic models tested (divergence plus migration). Pairwise migration rates among populations improved the likelihood of the divergence model, and the addition of further inter-population migrations highlighted that the most likely model is an asymmetric full island, although this model supports very little gene flow among these major worldwide populations (consistently ≪ 0.001 of effective population size per generation; Table 2B). The corrected AIC takes into account both number of parameters and number of observations, allowing for a consideration of differences in the likelihood comparison (Table S2). We ran these demographic inferences with and without redundant (near-identical) genome sequences from populations Africa2 and Africa1 (30, 31, 48 and 53) in order to correct for potential sampling bias, and obtained highly similar results.

Comparing population parameters estimated with different models indicates that the introduction of migration primarily influences results concerning the time of population splits and mutation rate (Table 2). While effective past and current population sizes have different absolute values, trends of population reduction (African populations) and growth (non-African populations) are confirmed throughout different models.

The timing of the two population splits, T2 and T4 (Figure 2, Table 2A), which presumably correspond to the out-of-Africa and American colonization events, are comparable to human estimates of population splits. Indeed, the second event appears to be 2 to 4 times more recent than the first (on average, ∼38k generations versus ∼110k generations, respectively), as expected under a bacterial-host model of co-expansion. According to models without migration, the estimate of divergence in number of generations of the Africa2 population from the other African strains (T1) also fits the timing of the divergence of the San population from other Africans, being twice as old as the out-of-Africa divergence (∼249k generations ago; Table 2A). Indeed, previous inferences based on human genetic data have estimated these events to have happened ∼60kya for the out-of-Africa (Eriksson et al. 2012), ∼20kya for the arrival into the Americas (Eriksson et al. 2012), and ∼110kya for San divergence (Veeramah et al. 2011; Hammer et al. 2011; Schlebush et al. 2012). On the other hand, the time inferred from the *H. pylori* dataset for the San split under the most likely model, which includes migration, is older than ∼500k generations.

**Figure 2.**
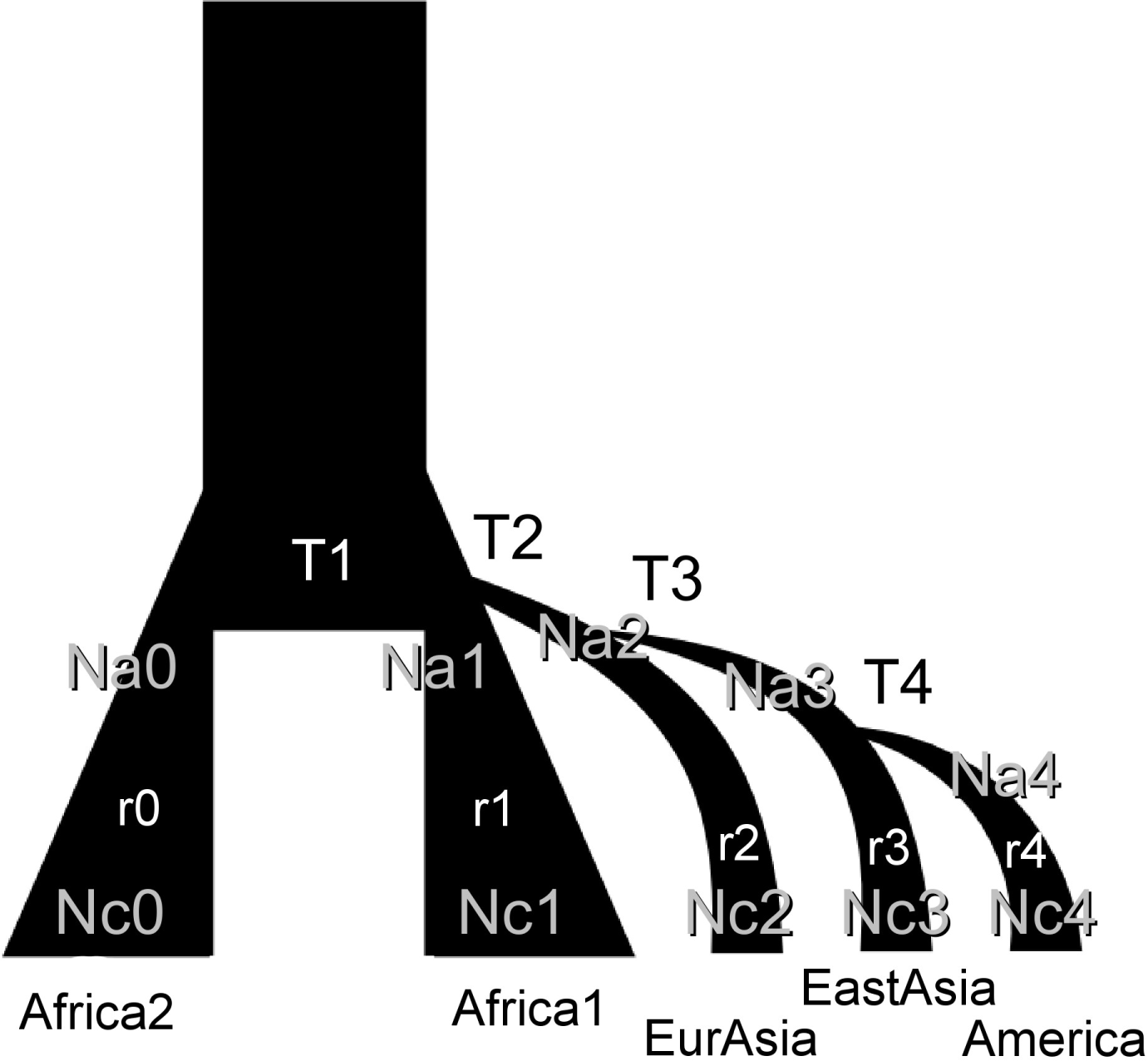
Schematic representation of the most likely genealogy inferred for *H. pylori* world-wide sample. Demographic parameters estimated via coalescent simulations are summarized. *T* parameters correspond to time of population splits (1 to 4, most ancient to most recent). *N*_*a*_ and *N*_*c*_ parameters indicate effective ancestral and current population sizes, with 0 being the Africa2 population and 5 the America population (most ancient to most recent). *R* parameters refer to population growth.

**Table 2. A).** Most likely demographic parameters estimated with *fastsimcoal2.1* for tree topology 2 and relative confidence intervals calculated for the migration model, which is the most supported. Parameters are reported assuming 2 generations/year. Population parameters correspond to those depicted in Figure 3; *r* parameters are the population growth rates, with the numeric order indicating populations from Africa2 to America (see Figure 3); *μ* is the mutation rate.

| Parameters |  | Ancestral effective populations size (Na) |  |  |  |  | Current effective population size (Nc) |  |  |  |  |
| --- | --- | --- | --- | --- | --- | --- | --- | --- | --- | --- | --- |
| Model |  | Na0 | Na1 | Na2 | Na3 | Na4 | Nc0 | Nc1 | Nc2 | Nc3 | Nc4 |
| No migration | Three pops | 1,918,265 | 309,326 | 171,636 | - | - | 106,580 | 898,799 | 3,178,763 | - | - |
| “ | Four pops | 1,852,974 | 809,566 | 477,655 | 51,188 | - | 101,030 | 230,367 | 2,307,144 | 1,875,418 | - |
| “ | Five pops | 2,373,071 | 1,083,279 | 293,795 | 28,444 | 18,449 | 103,453 | 257,756 | 2,015,267 | 761,583 | 690,042 |
| Migration | “ | 1,215,564 | 223,068 | 10,366 | 11,580 | 11,821 | 102,164 | 106,142 | 813,228 | 399,904 | 184,468 |
| c.i. 0.05 |  | 240,056 | 12,066.4 | 22,252.0 | 12,017 | 11,990 | 105,459 | 103,772 | 230,774 | 108,383 | 128,091 |
| c.i. 0.95 |  | 2,056,863 | 395,493 | 452,400 | 357,243 | 383,897 | 1,154,059 | 841,312 | 1,375,820 | 817,227 | 789,851 |

| Population splitting times |  |  |  | Population growth rates |  |  |  |  | Mutation rate |
| --- | --- | --- | --- | --- | --- | --- | --- | --- | --- |
| T1 | T2 | T3 | T4 | r0 | r1 | r2 | r3 | r4 | $\mu$ |
| 273,339 | 138,190 | - | - | 1.057e-05 | -3.902e-06 | -2.112e-05 | - | - | 8.069e-06 |
| 229,697 | 119,441 | 75,369 | - | 1.267e-05 | 5.472e-06 | -1.318e-05 | -4.778e-05 | - | 4.596e-06 |
| 245,942 | 128,714 | 45,670 | 31,778 | 1.274e-05 | 5.837e-06 | -1.496e-05 | -7.198e-05 | -0.000113 | 1.479e-07 |
| 529,626 | 89,686 | 69,096 | 44,338 | 4.675e-06 | 1.402e-06 | -4.864e-05 | -5.126e-05 | -6.196e-05 | 0.0009732 |
| 102,889 | 53,942 | 34,197 | 11,430 | -6.223e-06 | -9.437e-06 | -3.978e-06 | -7.891e-05 | -1.693e-04 | 0.0002284 |
| 350,810 | 95,913 | 51,312 | 28,388 | 1.434e-05 | 5.251e-06 | -3.558e-05 | 7.689e-06 | 1.172e-05 | 0.0008439 |

The long term mutation rate per site per generation estimated with *fastsimcoal2.1* varies between ∼8.47 × 10^−7^ and ∼9.73 × 10^−4^ (Table 2), this second estimate being much faster than the previous long term estimate, per site per year, from Morelli et al. (2010), based on the coalescent tree of the 7 housekeeping genes and inferred with ClonalFrame (2.6 × 10^−7^). Other previous estimates based on 78 gene fragments from serial and family isolates (1.4-4.5×10^−6^; Morelli et al. 2010), upon genomes sequentially taken from patients with chronic infection (2.5x10^−5^; Kennemann et al. 2011) and on genomes from 40 family members (1.38x10^−5^; Didelot et al 2013) are compatible with that inferred here by a purely divergent model. The bacterial recombination rate per initiation site per year obtained from our genomes analyzed with ClonalFrame (9.09 × 10^−9^) is more than 20 times slower than a previous estimate of 2.4 × 10^−7^ reported in Morelli et al. (2010), based on housekeeping genes using the same approach. It is important to note, however, that the recombination rate was not included in our models and that our absolute estimates are in generations instead of years.

Growth rates (*r*, see Table 2A) were negative for African clusters indicating population size reductions, with current effective population sizes (*N*_*c*_) being several times lower than ancestral population sizes (*N*_*a*_) for Africa2 and Africa1, respectively (Table 2A). The other three populations show signatures of expansion and appear to have been founded by a comparable few individuals, subsequently undergoing rapid growth. Migration rates are similarly small among pairwise populations, however outgoing migration rates from Africa are lower than the others (Table 2B). This result may indicate that gene flow did not extensively involve geographic macroareas, but if it did occur, mixed stains are more likely to be found in specific contact regions (e.g., coastal areas). Confidence intervals of demographic estimates with migration obtained using parametric bootstrap are reported in Table 2 and show important uncertainty associated with the best estimates.

**Table 2. B).** All *M* are pairwise migration rates numbered from population 0 (Africa2) to population 4 (America).

| **M01** | **M10** | **M12** | **M21** | **M23** | **M32** | **M34** | **M43** | **M02** | **M20** |
| --- | --- | --- | --- | --- | --- | --- | --- | --- | --- |
| 2.1092e-07 | 2.3405e-06 | 6.8818e-07 | 1.0032e-05 | 3.7624e-06 | 2.0730e-06 | 3.9028e-06 | 4.6873e-06 | 1.0517e-06 | 1.6399e-06 |
| 4.0924e-07 | 1.3297e-07 | 9.7725e-07 | 1.0824e-06 | 5.4905e-07 | 1.6803e-06 | 1.5580e-06 | 1.3040e-06 | 2.6228e-07 | 1.9271e-07 |
| 7.7355e-06 | 1.2793e-05 | 8.4155e-06 | 1.1639e-05 | 9.4577e-06 | 9.0864e-06 | 8.1415e-06 | 8.7104e-06 | 7.0558e-06 | 1.1664e-05 |

| **M03** | **M30** | **M04** | **M40** | **M13** | **M31** | **M14** | **M41** | **M24** | **M42** |
| --- | --- | --- | --- | --- | --- | --- | --- | --- | --- |
| 1.4843e-07 | 1.1139e-06 | 1.3739e-06 | 2.9639e-07 | 3.4169e-07 | 1.4061e-07 | 2.6611e-07 | 3.3381e-07 | 3.7917e-06 | 2.7227e-06 |
| 8.7827e-07 | 1.6083e-07 | 4.3319e-07 | 1.2945e-06 | 4.1117e-07 | 1.0548e-06 | 7.6409e-07 | 1.7891e-06 | 1.3625e-06 | 5.1306e-07 |
| 8.5424e-06 | 1.3895e-05 | 8.1077e-06 | 8.6616e-06 | 8.5431e-06 | 7.7131e-06 | 8.0440e-06 | 8.6578e-06 | 8.6370e-06 | 8.8781e-06 |

### Tests of positive selection and identified candidate regions

After correction of likelihood values with demographic simulations, the SweeD test of selection did not identify any strongly selected loci at the global level (Figure 3), but did indicate differential signatures of positive selection at the population level (Figure 3; Table 3). The largest number of selected loci was detected among African bacterial strains associated with San-speaking people (Africa2). Signatures of local positive selection were also observed in the Africa1 and American populations (Figure 3), while remaining populations (Eurasian and East Asian) did not show strong evidence for recent local adaptation (Figure 3).

**Figure 3.**
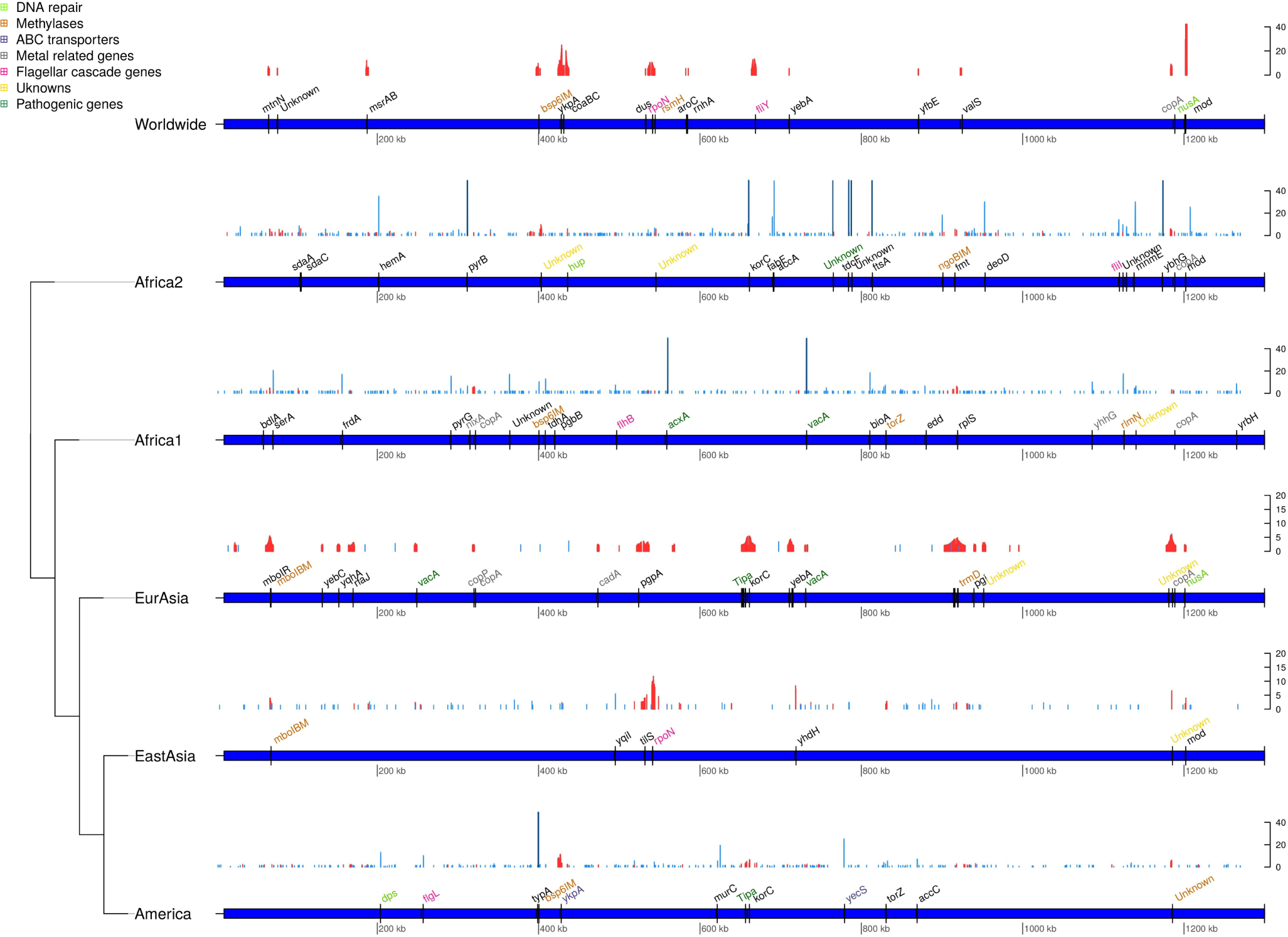
Results of the SweeD and OmegaPlus analyses. A comparative representation for a “synthetic” strain of the worldwide sample and one “synthetic” strain of each population is drawn using a fictitious topology. Selection values are reported on the graph above each synthetic strain, on the y-axis, and genomic position on the x-axis. Omega values are represented with red lines, while alpha values are reported in blue. Since alpha values reach much higher levels than omega values, to make the figure easy to read, we reported both omega and alpha values within a scale from zero to 50, and we indicated alpha values higher than 50 in darker blue. Genes falling into the functional categories explained in the discussion are color-coded as reported in the legend, while remaining are in black.

**Table 3.**
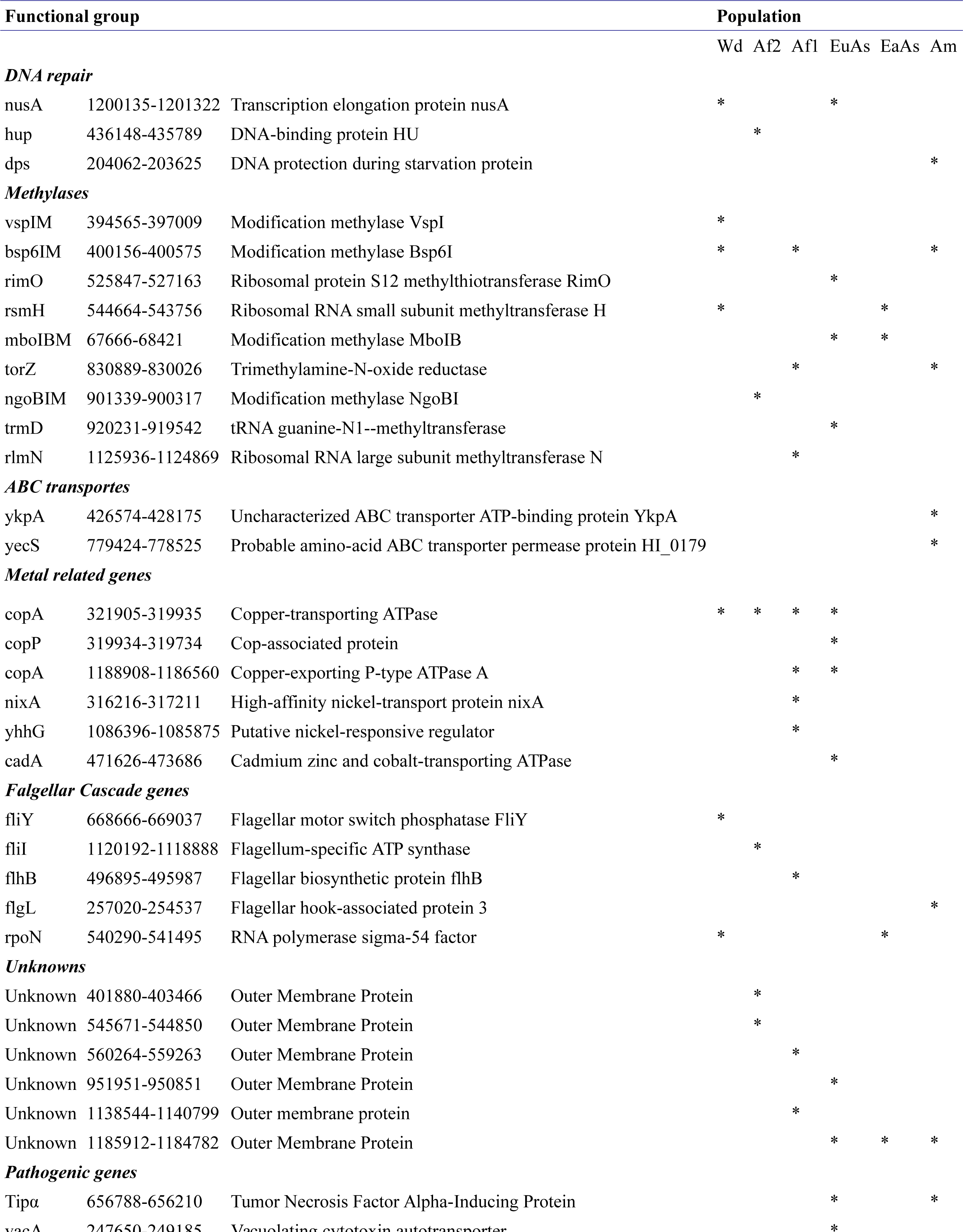
List of genes identified as being under positive selection by population, classified by function. Populations are abbreviated as Wd = worldwide; Af2 = Africa2; Af1 = Africa1; EuAs = EuroAsia; Eas = EastAsia; Am = America. For a complete list of genes identified as being putatively positively selected in worldwide and local samples see Table S3.

The same dataset analyzed with OmegaPlus, using as a null distribution the same demographic simulations analysed with SweeD, gave different results, with significance found mainly in the worldwide sample (Figure 3). The highest values of linkage disequilibrium were found in the global dataset (Table S3), with the highest peak associated with a gene coding for the elongation protein NusA, which has been studied in *Escherichia coli* (Cohen et al. 2010). Despite the structured nature of the worldwide sample, previous studies have demonstrated that population structure has little to no impact on the specific LD structure captured by the Omega statistic (e.g., Jensen et al. 2007a).

Both methods, SweeD and OmegaPlus, indicate several signatures of positive selection in African and American populations, while much lower signals are observed for Euro-Asian populations. The synthesis of the two analyses is presented in Figure 3. Regions that were significant for only one of the two methods were considered if their likelihood or omega value overcame the maximum value found for overlapping regions.

Using this approach, 158 genes are identified as putatively positively selected in either the total worldwide datasets or in the 5 sub-populations (Table S3 and S4), with the highest number (51) found in the Africa2 population. Moreover, this includes several unknown genes, most of which appear to code for outer membrane proteins (Table S4). Copper-associated genes (2 *copA* and 1 *copP*) are also indicated as positively selected. These genes are part of the *sro* bacterial operon and may relieve copper toxicity (Table S3; Beier et al. 1997; Festa and Thiele 2012). Among Africa2 strains, the highest likelihood values among Africa2 strains correspond to a well-known division protein gene (*ftsA*) (Figure 3 and Table S4). Moreover, the *pyrB* gene coding for aspartate carbamoyltransferase is also identified and was previously suggested as essential for bacterial survival (Burns et al. 2000). In the Africa1 population, the most important signal of selection appears associated to a *vacA* gene, a trait which has been consistently studied given its role in *H. pylori* pathogenic process (*e.g*. Basso et al. 2008; Yamaoka 2010). Other *vacA* and *vacA*-like genes are indicated in Africa2 and EuroAsian populations (Table S3 and S4).

## Discussion

Our analysis of a global *H. pylori* genome sample sought to illuminate both the selective and demographic histories of this human pathogen. Our analyses of population structure were carried out with particular attention, as population genetic clusters were the basic unit for demographic and selection inferences. Previous work based on MLST sequences and STRUCTURE software found a higher number of clusters distributed worldwide, a result largely accepted in the field. However, given the importance of population structure and the theoretical and computational limitations of some approaches, as well as the clonal reproductive behaviour of our organism, we explored population structure from complementary points of view (*i.e*. multivariate analysis, Bayesian analysis, co-ancestry analysis and coalescent genealogy). This combination of multiple approaches identified fewer populations globally, and thus offers an alternative perspective to previous results. Furthermore, our inferred mutation rate represents the first attempt to study the long-term substitution rate of *H. pylori* on a worldwide genome sample. Under a purely divergent model, the result was similar to the long-term rate previously estimated from MLST housekeeping genes (Morelli et al. 2010), but introducing migration led to much higher estimates.

While this analysis based on high-resolution data provides a reliable relative estimate of times to population divergence events, the open question remains on how to interpret and compare the bacterial inferences with those based on human genetics. Times of population splits T1, T2 and T4 are, in terms of the number of generations, roughly twice as old as has been proposed in the human demographic literature. If we use these estimates as calibration points to translate number of generations into years, we can deduce a number of bacterial generations per year = 2. An exception is represented by the estimate of San bacterial divergence when migration is accounted for, as the number of generations doubles to ∼530k translating into ∼265kya of split (still assuming a bacterial number of 2 generations per year). Notably, one recent estimate of San divergence obtained by Excoffier et al. (2013) is very near our estimate, *i.e*.∼260kya. If we alternatively used the latter estimate of split of Africa2 strains from others as a calibration point to deduce the number of bacterial generations per year, then we would consider that ∼530k bacterial generations happened within ∼110kya (which is the most supported estimate of San split from human genetic data). In this case, the number of generations per year would be ∼4.8 and the other times to bacterial population splits (T2, T3 and T4) would translate into much more recent events, although the relative timing of colonization of different geographic regions in absolute number of generations would not be affected.

*H. pylori* generation time is thus a key parameter in the estimation of co-evolutionary times of host-parasite population differentiation and also to make a comparison between our inferred long-term mutation rate with previous estimates which are calibrated in years instead of generations. Although two generations per year may seem unreasonably slow for a bacterial organism, we cannot exclude that the peculiar epidemiological dynamics of this bacterium, such as lifelong infection and acquisition early in life (see Introduction), may influence the long term generation time here considered. Both experimental (*i.e*. familial studies of age structured host samples) and analytical epidemiological models could be used to obtain an empirical estimate. Since *H. pylori* strains could not have colonized any area before the arrival of their human host, our proposed generational time can be considered a lower limit.

Apart from methodological limitations, the events and their timings elucidated here are largely congruent with the human genetic and archaeological literature, confirming previous hypotheses of a close co-evolutionary relationship between the two species (Linz et al. 2007; Moodley et al. 2012). The divergence of the African strains associated with the San, assuming a good fit between human and bacterial estimates, supports an ancient origin of human *Helicobacter* - seeming to have been already in association with the human host before the separation of the San population, and older than an association of at least ∼100 kyr suggested by MLST sequences (Moodley et al. 2012). Given the high level of host-specialization, one may hypothesize that this stomach pathogen evolved along with the human host early in the genus *Homo* – a model of interest for future investigation.

Most interestingly, from the bacterial perspective, are the strong signals of population size reduction within Africa, particularly dramatic in the case of the San-associated Africa2. This could have resulted from a reduction in the effective size of the human host population itself, as we know that San hunter-gatherer populations were adversely affected by the Bantu expansion (over 1000 years ago) and by more recent European colonization. However, this does not explain a similar but not as strongly negative growth rate in Africa1 strains, associated with the Bantu and other African populations, which are known to have increased in population effective size since the Neolithic revolution. One alternative to human demography may be stronger selection in Africa, a notion that is consistent with the larger number of putatively adaptive regions identified in Africa, relative to other sampled populations (Figure 3 and Table S3). Despite the very high prevalence of *H. pylori* on this continent, a significant association with the incidence of gastric diseases has never been demonstrated (Bauer and Meyer 2011; Graham et al. 2007). The opposite is true in non-African strains, where we show that *H. pylori* had a very low ancestral effective population size, coupled with the high population growth rates in our global sample. It may, therefore, be reasonable to hypothesize that the long-term African association of this bacterium with human populations may have led to selection for reduced pathogenicity, whereas a founder effect and rapid growth during the colonization of populations in other areas of the world could have freed this population from these long term selective constraints, possibly resulting in a more virulent and pathogenic bacterial population (Argent et al. 2008; Duncan et al. 2013). Concerning the divergence of the American population, we did not detect a clear signature of a founder event. Although the timing of the population split fits with the estimated human colonization of the Americas, we acknowledge an important lack of sampling coverage of the vast Siberia region which hinders more conclusive results on the expansion dynamics of *H. pylori* across East Asia and to the Americas.

The results obtained with different selection methods address somewhat different biological questions and the extent to which each of these is robust to non-equilibrium demographic histories has only been partially described (*e.g*., Crisci et al. 2013). Based on our inferred demographic history, however, it is possible to describe the true and false positive rates of these statistics for our specific model of interest – representing an empirical solution that may partially overcome such limitations. Global signatures of selection were found in association with several genes of unknown function.

### Worldwide and population-specific genes under selection

Patterns of local adaptation are potentially of great medical interest, as they may help explain the continentally-differing patterns of virulence observed thus far (Wroblewski et al. 2010; Bauer and Meyer 2011; Matsunari et al. 2012; Shiota et al. 2013). TheAfrica2 population shows the strongest evidence of recurrent local adaptation, a result which is perhaps intuitive given its long association with the San, one of the most ancient of human groups. Adaptive events within Africa2 include the protein coding *ftsA* gene (Table S3), which is associated with the cytoskeletal assembly during bacterial cell division (Loose and Mitchison 2014). In addition, results from the analysis of the Africa1 population highlight potentially interesting aspects of the long-term adaptation of *H. pylori* to this population.

Among European strains, we identified the only instance in which an antibiotic-associated gene (the penicillin binding protein 1A, *mrcA*; Table S3) was under selection. This gene was experimentally shown to confer resistance to β-Lactam when a single amino acid substitution occurs (Ser414→Arg; Gerrits et al. 2002). Although our annotated genome of the EuroAsian strains does not show this specific alteration, European *H. pylori* has been more likely exposed to antibiotic treatments than in other regions of the world. On the other hand, recent positive selection at the global level as a consequence of the use of antibiotics seems unlikely, as antibiotic treatment has not been implemented on a global scale. Surprisingly, our analysis did not detect relevant signatures of selection among EastAsian strains, despite the well-known medical risk of gastric cancer associated with these strains. The American population showed the strongest signature of selection associated with a GTP binding protein whose role is still unknown (*typA;* Table S3). Our overall results concerning putatively positively selected genes support the role of important metabolic pathways associated with structural and motility functions. This study thus highlights important candidates for future experimental and functional selection studies (for a complete list of candidate genes see Table S4).

#### Genes involved in DNA repair

Worldwide genomic regions under selection were identified by OmegaPlus (Table S3), with the strongest signature of selection at the transcription elongation factor gene *nusA*, also flagged locally among EuroAsian strains. In *Eschierichia coli*, this protein plays an important role in DNA repair and damage tolerance (Cohen et al. 2010). Since *H. pylori* infection of human stomachs can compromise host-cell integrity, inducing breaks in the double-strand and a subsequent DNA damage response (Toller et al. 2011), an efficient DNA repair mechanism could be important in protecting bacterial DNA from damage induced by itself or in response to altered physiological conditions in the host stomach. Along with this, indications of positive selection for genes protecting DNA integrity were found among Africa2 and American strains: HU binding protein (*hup*) and during starvation protein (*dps*), respectively (Table 3). The former protein protects DNA from stress damage in *H. pylori* (Wang et al. 2012), while the latter is required for survival during acid stress, although its role has been characterized in *E.* c*oli* but not in *H. pylori* (Jeong et al. 2008).

#### Genes involved in methylation patterns

Several genes expressing proteins involved in DNA methylation were identified as likely under selection (Table 3). A recent study by Furuta et al. (2014) used a genomic approach to compare methylation profiles of closely related *H. pylori* strains and showed outstanding diversity of methylation sequence-specificity across lineages. As methylation is an epigenetic mechanism responsible for the regulation of gene expression and phenotypic plasticity, the identification of certain selected methylation genes encourage the study of their specific role and their evolutionary implications in *H. pylori* methylation patterns.

#### ABC transporters

The ATP binding cassette (ABC) transporters are ubiquitous, and among their functions is the ability to expel cytotoxic molecules out of the cell, conferring resistance to drugs (Linton 2007). Two of these uncharacterised genes were indicated to be under positive selection in American strains (*ykpA* and *yecS*; Table 3).

#### Genes involved into flagellar cascade

Cell motility and cell adherence to the stomach mucosa is a key factors for the successful colonization of the human stomach, and several positively selected flagellum-specific genes (*flgL, flhB, fliI* and *fliY*) were identified across different local populations (Table 3).

Apart from genes involved into the flagellar cascade, positive selection was also detected in the regulating factor of the cascade itself (sigma(54) or *rpoN*), corroborating the importance of bacterial motility in survival (Table 3).

*Genes involved in heavy metal metabolism*. Importantly, our selection analysis highlights a potentially predominant role for genes associated with copper metabolism in the *H. pylori* life cycle, with the same genes flagged in multiple populations (Table 3). Copper mediated colonization of the stomach mucosa occurs through the action of trefoil peptides in *H. pylori* (Montefusco et al. 2013) and copper drastically increases in cancerous tissues. However the detailed role of *copA* and *copP* genes and of copper metabolism in *H. pylori* long-term adaptation is yet to be investigated. Interestingly, the Africa1 population shows signatures of positive selection of two genes involved in the transport and regulation of nickel (*nixA, yhhG*), while EuroAsian strains show hints of selection for a cadium, zinc and cobalt transporters (*cadA*; see Table 3).

#### Genes involved in virulence

We identified a number of putatively selected *vacA* genes in local populations as expected from previous indications of their importance in *H. pylori* pathogenicity (Olbermann et al 2010). It is further interesting to note that *vacA* and *vacA-like* genes also show evidence for selection among African populations, where the association of *H. pylori* with gastric disease is not considered to be significant. In particular, Africa1 strains present a strong signal associated with the acetone carboxylase beta subunit (*acxA*; Table 3), which is part of the pathologically relevant operon *acxABC*, as it is associated with virulence and survival of the bacterium into the host stomach (Brahmachary et al. 2008; Harvey 2012). These observations suggest that virulence-related genes may nonetheless play an important role in bacterial adaptation or, more specifically, that *H. pylori* may indeed have a pathogenic role among African populations that is masked by other factors leading to gastric diseases. Finally, the Tumor Necrosis Factor Alpha-Inducing Protein (*Tipα*; Suganuma et al. 2001; 2006; 2008) was identified in EuroAsian and American strains, calling for closer investigation in relation to its potentially pathogenic role among these specific populations.

#### Outer membrane proteins (OMP)

Many unknown genes appear into the list of putatively selected genes (Table 3). Among those, there could be a particular interest in further investigating the nature and role of outer membrane proteins, which would certainly provide valuable information on the interaction of *H. pylori* and the gastric environment. There are at least five recognized families of genes coding for OMP (HopA-E), which are involved in the processes of adherence to the gastric mucosa and thus play an important role in successful colonization of the host's stomach (Oleastro and Ménard 2013; Yamaoka and Alm 2008). Moreover, the importance of specific OMP genes in *H. pylori* has been investigated in recent studies (Kennemann et al. 2012; Nell et al. 2014).

From an evolutionary perspective, our study presents evidence for processes of adaptation in *H. pylori* to its human host, but, regrettably, does not provide a perspective on the co-evolutionary interactions that are likely to have occurred during their long history of association. In this sense, it is intriguing to speculate that the interaction with the human host did not simply lead to pathogenic conditions but also to mutual adaptation. Theories on beneficial interactions of *H. pylori* and the human host have been already suggested (Blaser 2008). The observation that fewer than 15% of infected human individuals show clinical symptoms has led previous studies to speculate that *H. pylori* may play an important, but not necessarily pathogenic, role in the human gastric niche, potentially even protecting its host from other gastric infections (Shahabi et al. 2008; Blaser 2008; Atherton and Blaser 2009). In support of this idea, a recent survey among native Americans reported that patients with lower host-bacteria co-ancestry - that is, patients infected with hpEurope (here included into the Eurasian population) and not with hspAmerind (the American population) - show increased severity of premalignant lesions in gastric cancer (Kodaman et al. 2014). Hopefully, future investigation will also focus on the long-term interaction of the two species and the possible signatures in the human genome that result from the long association with *H. pylori*.

Although our results highlighting major selective events in Africa are supported by a common African origin for both species, the co-evolutionary history between *H. pylori* and humans is an area that warrants future and more detailed investigation at the genomic level. A first step would be the inclusion of more genomes from underrepresented regions such as Sahul, North-East Africa, Central Asia and the Americas. Furthermore, unrepresented regions such as Siberia and Oceania would allow for the investigation of genetic continuity/discontinuity across north-eastern and south-eastern Asia to the Americas and the Pacific, respectively. A deeper analysis of Asian, American and Austronesian bacterial genomes may also help shed light on alternative Pacific routes for the colonization of the Americas, a hypothesis that has been widely debated in the literature (see Gonçalves et al. 2013; Malaspinas et al., 2014).

## Acknowledgements

This project was supported by ERA-NET PathoGenoMics project HELDIVNET (0313930B) from the German Ministry of Education and Research (BMBF) to SS and BMBFproject 01GS0805 to RR for massive parallel sequencing. VM was supported by a postdoctoral fellowship from the European Union (Framework 7) and a short term post-doctoral fellowship from the European Molecular Biology Organization (EMBO). JDJ and MF were supported by grants from the Swiss National Science Foundation and a European Research Council (ERC) Starting Grant to JDJ. XD would like to acknowledge the NIHR for Health Protection Research Unit funding. We would like to thank the Vital-IT bioinformatic center for the technical support with the Vital-IT cluster. We also thank Mark Achtman for useful comments on an early version of the manuscript.

